# “The School Cafeteria Problem”: Disrupted Visuospatial Attention During Multisensory Speech-in-Noise Perception in Children with ADHD

**DOI:** 10.1101/2025.11.11.687600

**Authors:** Nishant J. Dinesha, Sarah Mehta, Zakilya Brown, Alesia A. Richardson, Rachel A. Rivera, Christina Layton, John J. Foxe, Emily J. Knight

## Abstract

**Background:** Many children with attention deficit hyperactivity disorder (ADHD) have difficulty distinguishing speech in crowded scenarios with competing conversations and noise. This can impact learning and community participation. Prior work has implicated audiovisual integration of lip movements with speech sounds as a potential underlying mechanism, but difficulty in accurately deploying spatial attention is an important, often overlooked, factor that may impact the accuracy of speech perception in noisy environments.

**Objective:** Evaluate patterns of audiovisual integration and allocation of visuospatial attention during speech processing in a multiple-talker scenario among school-age children with ADHD and typically developing children (TD).

**Methods:** We recruited 7–12-year-old children with ADHD to participate in an integrated virtual reality and electrophysiologic (EEG) speech-in-noise perception paradigm. In virtual reality, children were asked to press a button whenever a centrally-located target character spoke a target monosyllabic word, while ignoring any words spoken by two flanking distractor characters. We manipulated 1) audiovisual (AV) content by intermixing trials containing either audio, visual, or multisensory AV representations of the three characters and 2) spatial attention demands by including trials with and without interference from the distracting speakers. We measured AV gain in target detection accuracy. Simultaneously, steady-state visual evoked potentials (SSVEP) were continuously recorded via a 128-channel electrode array as an index of visuospatial attention allocation to each speaker. To elicit SSVEP, the target and distractor speakers were tagged with different visual frequency oscillations (23hz and 15hz, respectively). EEG signal-to-noise ratio (SNR) in the corresponding first and second harmonics for those frequency bands was measured over the occipital scalp.

**Results:** Children with ADHD showed reduced AV gain relative to TD controls. TD controls also demonstrated more robust SSVEP at 23Hz and 46Hz (target speaker) than at 15Hz (distractor speakers). Compared to TD children, children with ADHD had reduced selective activation in response to the target. As a result, children with ADHD showed more equivalent allocation of visuospatial attention between 23Hz target and 15Hz distractor frequencies.

**Conclusions:** Children with ADHD demonstrate a reduction in the typical perceptual benefit afforded by exposure to multisensory compared to unisensory speech stimuli, accompanied by substantial differences in the allocation of visuospatial attention to the relevant speaker. Thus, dysregulation of selective visuospatial attention may impair their ability to effectively perceive language in noisy settings.

## INTRODUCTION

Children with attention deficit hyperactivity disorder (ADHD) often have difficulty distinguishing speech in the setting of other distracting stimuli (i.e., speech-in-noise perception) (Blomberg et al., 2019). Parsing speech from background noise is essential to promote effective social communication and classroom-based learning, presenting a “school cafeteria problem” for children analogous to the “cocktail party scenario” that has received much attention in the adult auditory perception literature {Cherry, 1953 #263;Ross, 2022 #265}. Thus, these auditory processing disorders can have far-reaching negative psychosocial impacts (Kreisman et al., 2012).

Prevalence estimates suggest that up to 40% of children with ADHD and up to 75% of children with other developmental disabilities manifest auditory processing difficulties (Aristidou and Hohman, 2025; Dawes and Bishop, 2009; Kreisman et al., 2012), underscoring the broad public health impact of these symptoms. Yet, the field lacks effective evidence-based diagnosis and intervention strategies (Dawes and Bishop, 2009; McArthur, 2009). Interventions in current use that attempt to minimize sensory symptoms in developmental disabilities (e.g., frequency modulation systems, auditory integration training) were developed long before the initiation of systematic studies on auditory processing disorders, have unclear connections to the mechanisms uncovered in research, and have little empirical support (McArthur, 2009). It is critical to enhance our understanding of the specific mechanistic differences underlying auditory processing disorders in ADHD to support the design of evidence-based diagnosis and targeted intervention strategies.

Multiple processes have been implicated in the successful perception of speech in noisy environments. A substantial body of literature has focused on the role of automatic (pre-attentive) segmentation of auditory inputs into distinct streams in facilitating speech in noise perception (Calcus, 2024; Sussman, 2017). Selective auditory attention is thought to interact with this segmentation process to augment the perception of behaviorally relevant stimuli (Alain et al., 2001; Sussman, 2013, 2017). These processes appear to mature over early childhood, with school-age children exhibiting less efficiency in automatic auditory scene segmentation than young adults, requiring greater separation between competing streams (Sussman and Steinschneider, 2009; Sussman et al., 2007) and a higher dependence on auditory attention to promote accurate behavioral performance (Sussman and Steinschneider, 2009). Furthermore, children with developmental language disorders exhibited additional delays in the maturation of auditory stream segmentation relative to same chronologic age peers (Sussman et al., 2015).

However, auditory perception does not occur in isolation in real-world multi-talker environments. Audiovisual (AV) speech perception recruits both visual and auditory cortices, enhancing neural connectivity between these regions (Peelle et al., 2022). It is well known that visual information from a speaker’s lip movements helps disambiguate speech when acoustic signals are convoluted, highlighting the relevance of AV integration to speech perception in the context of noise (Iarocci et al., 2010; Ma et al., 2009; Ross et al., 2007a). Visual cues can provide complementary information to auditory inputs that help to predict the timing of auditory stimuli, enhance phoneme perception (Yuan et al., 2021), and serve as spatial cues to help differentiate speech streams during interference (Haykin and Chen, 2005). As a result, much prior research has focused on AV integration deficits as a potential driver for speech-in-noise perception problems in ADHD and related developmental disabilities (Foxe et al., 2020; Foxe et al., 2015; Gijbels et al., 2021; Irwin and DiBlasi, 2017; Lalonde and Werner, 2021; Ross et al., 2007b; van Laarhoven et al., 2018; Vanneau et al., 2025). Adults with ADHD have demonstrated reductions in AV benefits during speech-in-noise perception (Michalek et al., 2014; Schulze et al., 2021). They are also less affected by the McGurk illusion, in which incongruous lip movements influence the perception of speech syllables (Schulze et al., 2021). Likewise, EEG studies have shown alterations in event-related potentials across frontal, parietal, and occipital brain regions during AV integration tasks in young adults with ADHD (McCracken et al., 2020; McCracken et al., 2019). Some studies have also identified deficits in non-speech AV integration in ADHD, such as narrowed temporal binding windows (Panagiotidi et al., 2017). However, others suggest that such AV integration deficits are specific to the speech domain. For example, unlike their reduced susceptibility to the McGurk illusion, adults with ADHD have shown intact responsiveness to a non-speech AV illusion, the sound-induced flash illusion (Schulze et al., 2022).

However, as with auditory stream segmentation, AV integration may interact with attentional processes. Congruent visual inputs have been shown to enhance a listener’s ability to attend to the relevant speaker amidst competing auditory inputs and lead to diminished representation of auditory inputs from unattended speakers (Zion Golumbic et al., 2013). Thus, the possibility that deficits in visuospatial attention contribute to the observed deficits in AV integration and the resultant difficulty with speech-in-noise perception exhibited by children with ADHD is critical to explore. Indeed, studies suggest that children with ADHD may exhibit more substantial impairments in visual than in auditory attention performance (Lin et al., 2017). The orientation of visual attention to the face or speech sounds can also interfere with efficient AV integration in other related developmental disability populations, such as autism (Tietze et al., 2019), and some evidence suggests that children with ADHD have similar alterations in patterns of gaze fixation to the face (Frick et al., 2023; Lin et al., 2017). One study of adults with ADHD revealed multisensory impairment during high but not low perceptual load conditions (Schulze et al., 2022), lending additional support to the notion that attentional demands may influence AV integration in this population. Nevertheless, how interactions between AV integration and visuospatial attention processes contribute to speech-in-noise processing deficits in children with ADHD is not well understood.

A useful electrophysiologic index that has been employed to help assess the allocation of visuospatial attention toward competing stimuli is the steady-state visual evoked potential (SSVEP) (Senkowski et al., 2008). SSVEPs are neural responses evoked by oscillatory visual stimuli presented at specific frequencies, with increased EEG responses observed when participants direct attention to these stimuli (Yang et al., 2022). Studies have shown that SSVEP amplitudes are largest during focused attention, intermediate during divided attention, and lowest when unattended (Yang et al., 2022). SSVEPs are also correlated with alpha band activity, a well-known neurophysiologic indicator of attention (Davidson et al., 2020; O’Connell et al., 2009). This measure has previously been employed to assess visuospatial attention during AV speech-in-noise perception in typically developing (TD) adults. Individuals who performed well on the task demonstrated clear selective activity in their SSVEP to the target speaker. In contrast, highly distracted participants who responded to the off-target speaker at greater rates demonstrated less target specificity in their SSVEP (Senkowski et al., 2008). Thus, SSVEPs are a useful and behaviorally relevant marker of visual attention allocation during multiple-talker speech perception tasks. SSVEPs can also reliably classify the locus of attention with high accuracy in more complex tasks such as controlling avatars in virtual environments (Yang et al., 2022), opening up new, more naturalistic avenues for study. Prior literature has primarily relied on relatively simplistic stimulus pairings to explore AV integration. In the natural world, stimuli are considerably more complex and exhibit systematic congruencies whereby visual features interact with auditory signatures (Besson et al., 2011). Our extensive experience as humans with these natural stimuli fundamentally shapes our multisensory perception (Senna et al., 2021; Snyder et al., 2015; Stein et al., 2014; Wallace et al., 2004). Attention to sensory inputs also varies from the lab to real-world settings (Peelen and Kastner, 2014). Thus, moving toward more naturalistic paradigms that offer increased ecological validity is helpful in evaluating the role of AV integration and attention deficits in clinical auditory processing disorder symptoms.

Here, we applied a virtual reality (VR) adaptation of the paradigm developed by Senkowski and colleagues (Senkowski et al., 2008) to evaluate patterns of AV integration and allocation of visuospatial attention during speech-in-noise among school-age children with ADHD and a cohort of age-matched TD control children. Additionally, we investigated brain-behavior relationships between the neurophysiologic findings and clinical measures of disordered auditory processing. Overall, the results revealed disruptions in the allocation of visuospatial attention in children with ADHD, which were associated with reduced multisensory AV benefit and poorer target detection accuracy under a multiple competing speaker scenario.

## METHODS

### Participants

Thirty-six children with ADHD and thirty-three group age-matched TD children aged 7–12 years were initially enrolled in the study. Five children with ADHD were excluded from the analysis for poor data quality (defined as less than one experimental block/five minutes of SSVEP data remaining after exclusion of artifacts), two TD children were excluded for failing to complete all ten experimental blocks, and two TD children were excluded for new developmental diagnoses reported after enrollment. Therefore, final primary analyses were conducted on 31 ADHD and 29 TD children. Included participant characteristics are outlined in Table 1. One participant in the TD group reported selective serotonin reuptake inhibitor use for treatment of anxiety. Twenty-two participants in the ADHD group reported chronic therapy with one or more psychotropic medications (stimulants: n=17, alpha agonists: n=10, selective serotonin reuptake inhibitors: n=16, atomoxetine: n=1). For all patients taking psychotropic medications, the medication dosage was stable for at least one month. Participants were asked to hold stimulant medications on the day of electrophysiologic recording, when possible, due to the demonstrated impact of stimulant medications on evoked potentials (Sawada et al., 2010). Sixteen out of seventeen children who routinely take stimulants were able to comply with this request. With the exception of three TD and seven ADHD participants who were lost to follow-up, all children participated in neuropsychological phenotyping, including the Wechsler Intelligence Scale for Children-5^th^ Edition (Kaufman et al., 2015) SCAN-3:C Tests for Auditory Processing Disorders (Keith, 2009) and the abbreviated Swanson, Nolan, and Pelham (SNAP-IV) to measure ADHD symptoms (Hall et al., 2019). In addition, participants with ADHD were required to have a prior diagnosis of ADHD according to the Diagnostic and Statistical Manual-5^th^ Edition (DSM-5) criteria by a clinical provider (American Psychiatric Association, 2013). Participants who were lost to follow-up for cognitive testing are included in the behavioral and electrophysiologic analyses but excluded from the neuro-phenotypic correlational analyses. The University of Rochester Research Subjects Review Board approved all experimental procedures. Each participant provided written informed consent and children provided developmentally appropriate assent.

**Table 1.** Participant Characteristics.

|  | **ADHD (n=31)** | **TD (n=29)** | **Significance (p)** |
| --- | --- | --- | --- |
| **Age** | 9.4±1.7 | 9.2±1.5 | $p=.728$ |
| **Sex** | 68% Male | 52% Male | $p=.206$ |
| **WISC-V Full Scale IQ<sup>a</sup>** | 108±14.4 | 113±13.0 | $p=.148$ |
| **WISC-V Verbal Comprehension Index<sup>a</sup>** | 115±17.1 | 113±13.5 | $p=.644$ |
| **SNAP-IV Inattention** | 18.4±4.7 | 5.5±4.4 | $p=8.2 \times 10^{-16}$ |
| **SNAP-IV Hyperactivity and Impulsivity** | 16.3±5.3 | 3.5±3.4 | $p=6.9 \times 10^{-16}$ |

Note: Mean ± standard deviation is presented for age, IQ and SNAP-IV scores. The proportion of males is presented for sex. <sup>a</sup>IQ data not available for n= 7 ADHD and n=3 NT subjects.

### Stimuli

The VR stimulus was comprised of three photorealistic 3D virtual characters (one target and two distractor speakers), each 1.6m in height, positioned at a fixed location within a virtual room (18m × 20m × 14m). The characters were viewed from a distance of 2m, with 0.5m separating each character. The visual angle between adjacent characters (measured from mouth to mouth) was 14.25°, and the central character’s height subtended a visual angle of 43.6°. The characters uttered a subset of 10 consonant-vowel-consonant (CVC) monosyllabic words that were in the lexicon of children (bag, cat, dog, gum, hug, jam, mid, row, sun, toy) selected from a previously studied stimulus set (Ross et al., 2007a). MakeHuman™ 1.2.0, an open-source graphics middleware, was employed to prototype the 3D photorealistic virtual characters. NVIDIA Omniverse™ Audio2Face technology (Version 2022.2.1) was then utilized to generate facial motion and lip-sync animations, with a prediction delay of 0.1 seconds, corresponding to a 100ms visual lead relative to the trimmed (958ms duration) audio files of the monosyllabic words (See Figure 1a). Blender (Version 3.6.0-usd.200.1), an open-source graphics software, was used to render the facial lip-sync animations for the virtual characters at 25 frames per second, which were subsequently exported in an FBX file format to the cross-platform Unity game engine (Version 2021.3.24f1), which was employed for stimulus presentation via a Meta Quest 2 (Meta, USA) virtual headset. Quality of the lip-sync was verified in pre-experimental pilot testing.

**Figure 1.**
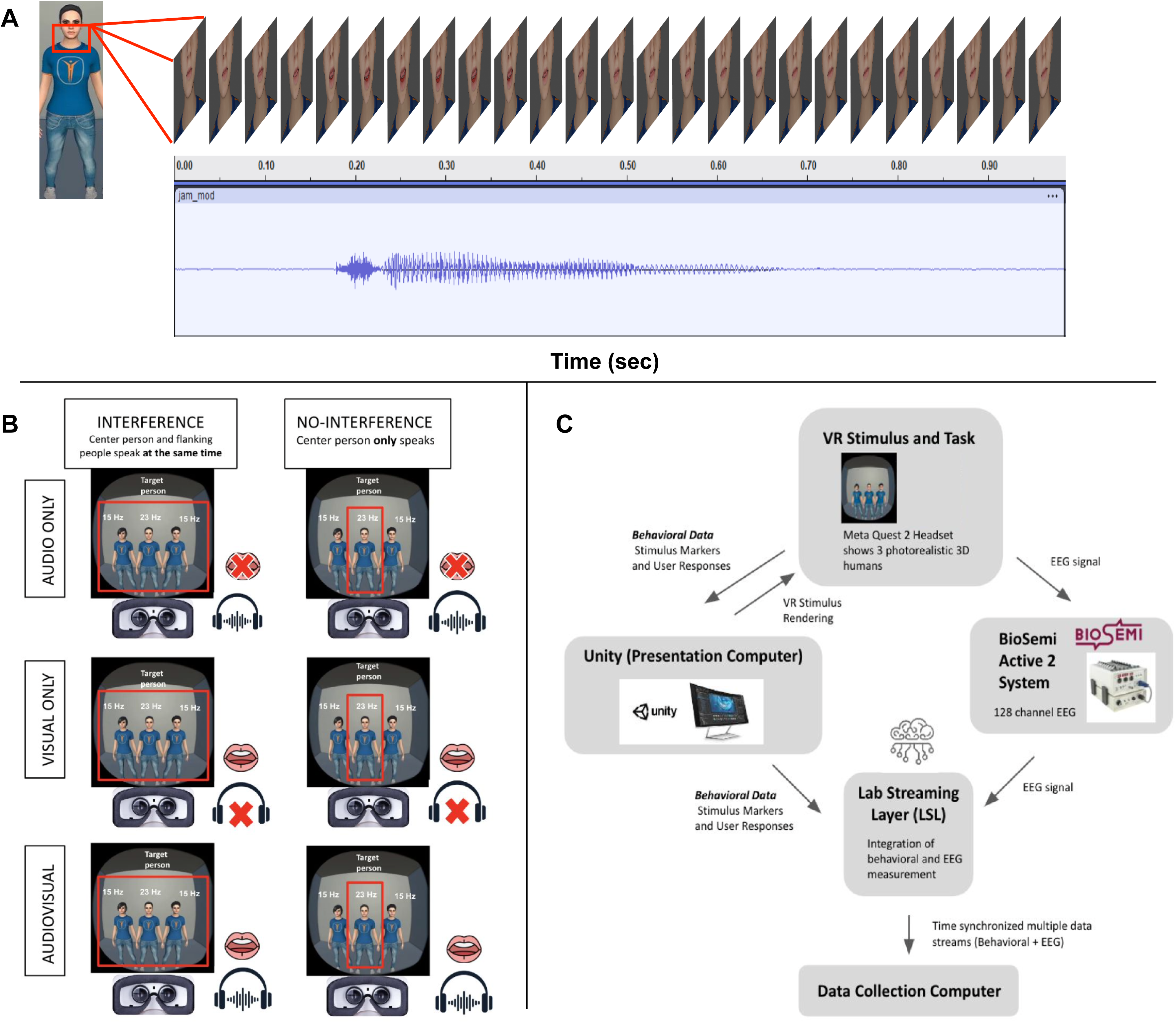
**A)** Schematic depiction of the onset and duration of visual and auditory stimulation in a representative audiovisual trial. Facial lip-sync animations are presented with a 100-millisecond visual lead relative to the trimmed (958ms duration) audio files of one of ten consonant-vowel-consonant monosyllabic words (i.e., bag, cat, dog, gum, hug, jam, mid, row, sun, toy). **B)** Schematic depiction of the two experimental manipulations (interference vs. non-interference and visual-only vs. audio-only vs. audiovisual) and the resultant six trial types. **C)** Schematic depiction of the experimental setup. Stimuli are rendered in Unity and presented via a Meta Quest 2 headset and headphones. Trial information, behavioral button press data, and electrophysiologic data are time-synchronized using Lab Streaming Layer (LSL).

The experiment consisted of randomly intermixed conditions (interference and non-interference) and sensory modalities: audio-only, visual-only, and audiovisual (AV) (See Figure 1B). In the interference condition, all three virtual characters concurrently uttered a CVC monosyllabic word; in the non-interference condition, only the central target speaker uttered a CVC monosyllabic word. In the AV condition, the auditory words were presented in conjunction with the corresponding visual facial lip-sync animations of virtual characters articulating the corresponding words; in the audio-only condition, the auditory words were presented in conjunction with still virtual characters (3 characters with no facial lip-sync animations). Finally, in the visual-only condition, only the visual facial lip-sync animation of virtual characters articulating the words was presented with no audio.

The VR stimulus presentation was divided into 10 blocks to mitigate prolonged continuous exposure to the VR headset and provide adequate breaks. The target word was varied between experimental blocks, and the order of blocks was counterbalanced across participants. Each block consisted of a total of 180 trials (2 conditions * 3 modalities * 30 trials), randomly presenting the above conditions and modalities intermixed with 12 (40%) target word trials in each category (conditions * modalities). Interstimulus interval (ISI) duration was randomly varied between 0 and 300ms with a mean of around 160ms. The monosyllabic auditory words were presented at approximately 74dBA sound pressure level (SPL) with a background pink noise around 59 dBA (SPL) over headphones (Sennheiser, model HD 600).

### SSVEP

To index the participants’ allocation of visual spatial attention towards three speakers, two blocks (first and last) had visual stimulation on a frame-by-frame basis as follows: the center speaker alternated between on and off periods every 21.71ms, which corresponded to a visual on-off flickering frequency of 23 Hz. The two distracting side speakers had visual stimulation with on-and-off periods every 33.30ms, corresponding to a flickering frequency of 15 Hz. The refresh rate of the Meta Quest 2 device was set to 72 Hz. Stimulus frequencies were selected such that their harmonics were non-overlapping with one another and with the refresh rate.

### Task

Participants were instructed to press a button on the Meta Quest 2 controller immediately upon hearing the target word from the center speaker while ignoring words from the distracting speakers. No feedback was provided regarding response accuracy. Prior to the main task of 10 blocks, all participants completed a practice block of 30 trials (2 conditions × 3 modalities × 5 trials), presented in randomized order and interspersed with 2 (40%) target word trials per category (the target word for the demo block was different from the main blocks). If a participant failed to respond to target words or demonstrated random button presses, the practice block was repeated, accompanied by a reiteration of the instructions. To enhance engagement, the practice block was introduced with a narrative involving a fictional friendly alien character requesting task completion according to the provided instructions.

### Data Acquisition and Preprocessing

EEG data were acquired using a BioSemi ActiveTwo system (Amsterdam, The Netherlands) with 128 scalp electrodes at a sampling rate of 512 Hz with the Common Mode Sense (CMS) and Driven Right Leg (DRL) electrodes as the reference. EEG data, stimulus presentation, and task event timing were synchronously streamed to a computer via LabStreamingLayer (LSL) (See Figure 1C). EEG data were processed offline using custom MATLAB (The MathWorks, Natick, MA, USA) scripts utilizing EEGLAB (EEGLAB v14.1.2b) (Delorme and Makeig, 2004) and ERPLAB toolboxes (Lopez-Calderon and Luck, 2014). EEG signals were filtered using an IIR Butterworth bandpass filter between 0.1 and 200 Hz implemented in ERPLAB. Filtered data were subjected to automated artifact rejection, including the removal of noisy channels (channel correlation criterion = 0.65; line noise criterion = 4) and high-amplitude artifacts (burst criterion = 30), using the pop_clean_artifacts function from the clean_rawdata plugin. Subsequently, spherical interpolation was applied to reconstruct the removed channels, followed by average re-referencing. This resulted in a median (interquartile range) acceptable recording duration of 7.9 (1.9) minutes and 8.2 (2.1) minutes for the ADHD and TD groups, respectively.

### Frequency Domain Analysis

To characterize the spectral features of the SSVEP blocks, the first and last blocks of pre-processed EEG data were concatenated and transformed into the frequency domain using MATLAB’s pwelch function with parameters set to N=1024 FFT points, a window length equal to N, and 50% overlap. Amplitude spectra (µV) for each channel were obtained by computing the square root of the pwelch output at each frequency bin (frequency resolution of 0.5 Hz). To enhance the detection of spectral peaks, the signal-to-noise ratio (SNR) was calculated at each frequency bin as the amplitude at that frequency bin divided by the mean amplitude in the surrounding 3 Hz frequency bins, excluding the adjacent bins (Liu-Shuang et al., 2014; Retter and Rossion, 2016; Rossion et al., 2012). Harmonic frequency responses (2F, 3F, etc.) were expected for each stimulating frequency (15 and 23 Hz) (Retter and Rossion, 2016). The number of significant harmonics included in further analysis was determined by computing grand-averaged SNR spectra across pooled groups (Typical, ADHD) and channels, and assessing significance via z scores (difference between the SNR at the frequency of interest and the mean SNR of the surrounding 3 Hz bins, divided by the standard deviation of those bins). Harmonics were included if their z scores exceeded 2.32 (p < 0.01, one-tailed) (Retter and Rossion, 2016). Only the distractor fundamental frequency of 15Hz and the target fundamental frequency of 23Hz, along with its second harmonic of 46 Hz (2*23Hz), satisfied the above criteria. For the target, we summed the SNR of 23 Hz and its second harmonics, 46Hz. Whereas for the distractor, we used only the SNR of 15Hz, as the second harmonic, 30Hz, did not satisfy the criteria. Considering each of the first and second harmonics separately (23Hz, 46Hz, 15Hz, and 30Hz) does not change the overall pattern of group effects (See Table S1). The electrodes of interest (EOI) were selected as the union of electrodes that showed the largest SNR responses in the pooled data (irrespective of group) (Rossion et al., 2015) to the target (23+46 Hz) and distractor (15 Hz) frequencies, which resulted in selection four midline channels around POz and Oz (see Figure 3C). For statistical analysis, the mean SNR of EOI channels for each participant was analyzed using mixed-design ANCOVA with age as a continuous covariate.

### Behavioral Analysis

The behavioral data were recorded via button presses across all 10 blocks, facilitated by the Unity Experiment Framework (Brookes et al., 2020). Data from all blocks were included to allow for sufficient behavioral data collection to be considered in the main analyses (percent correct and false alarms are also depicted separately by group and condition in Figure S1, demonstrating no difference in the pattern of behavioral results between flickering and non-flickering blocks). Target detection accuracy for each condition (interference and non-interference) and modality (AV, audio-only, and visual-only) was calculated using d-prime (d’). In signal detection theory, d’ serves as an index of the actual signal relative to the noise and can be measured as the difference between the normalized hit rates and false alarm rates (d’ = Z(hit rate) – Z(false alarm rate)). Corrections were applied to avoid undefined d’ values resulting from hit rates or false alarm rates of 0 or 1. (Hautus, 1995). Accuracy was then compared across between groups, conditions, and sensory modalities using a mixed design ANCOVA with age as a continuous covariate. Given that the primary metric of interest was AV gain, only AV vs audio-only performance was included in this analysis. However, percent correct and false alarms for each group during the visual-only condition are presented in Figure 1C for completeness and transparency.

## RESULTS

### Behavioral Performance

Target detection accuracy is depicted by group and condition in Figure 2, and the results of the statistical analysis are summarized in Table 2. There was a main effect of age on target detection accuracy, such that older children showed superior performance compared to younger children (F(1,57)=7.306, p=.009, η^2^_p_ =.114), irrespective of group. As expected, there was also a main effect of interference with better performance noted in the non-interference than interference condition (F(1,57)=12.108, p=.00097, η^2^_p_=.175). After covarying for age, TD chilldren displayed overall higher target detection accuracy across all conditions (F(1,57)=4.840, p=.032, η^2^_p_ =.078). Additionally, there was a significant group x sensory condition interaction (F(1,57)=5.028, p=.029, η^2^_p_=.081) with the TD group showing significantly higher audiovisual gain in accuracy relative to the ADHD group (Interference: AV Gain_TD_= .282 ± .398, AV Gain_ADHD_= .067 ± .412, t(58)=-2.052, p=.045; Non-interference: AV Gain = .501 ± .916; AV Gain_ADHD_=.101 ± .571F(1,56)=4.044, p=.049, η^2^_p_ =.067, t(58)=-2.043, p=.046). No other main effects or interactions reached significance. There was no substantial difference in the overall pattern of behavioral responses between flickering and non-flickering blocks (See Figure S1); thus, the stimulus flickering necessary to generate an SSVEP did not appear to artificially impact our measure of performance.

**Table 2.**
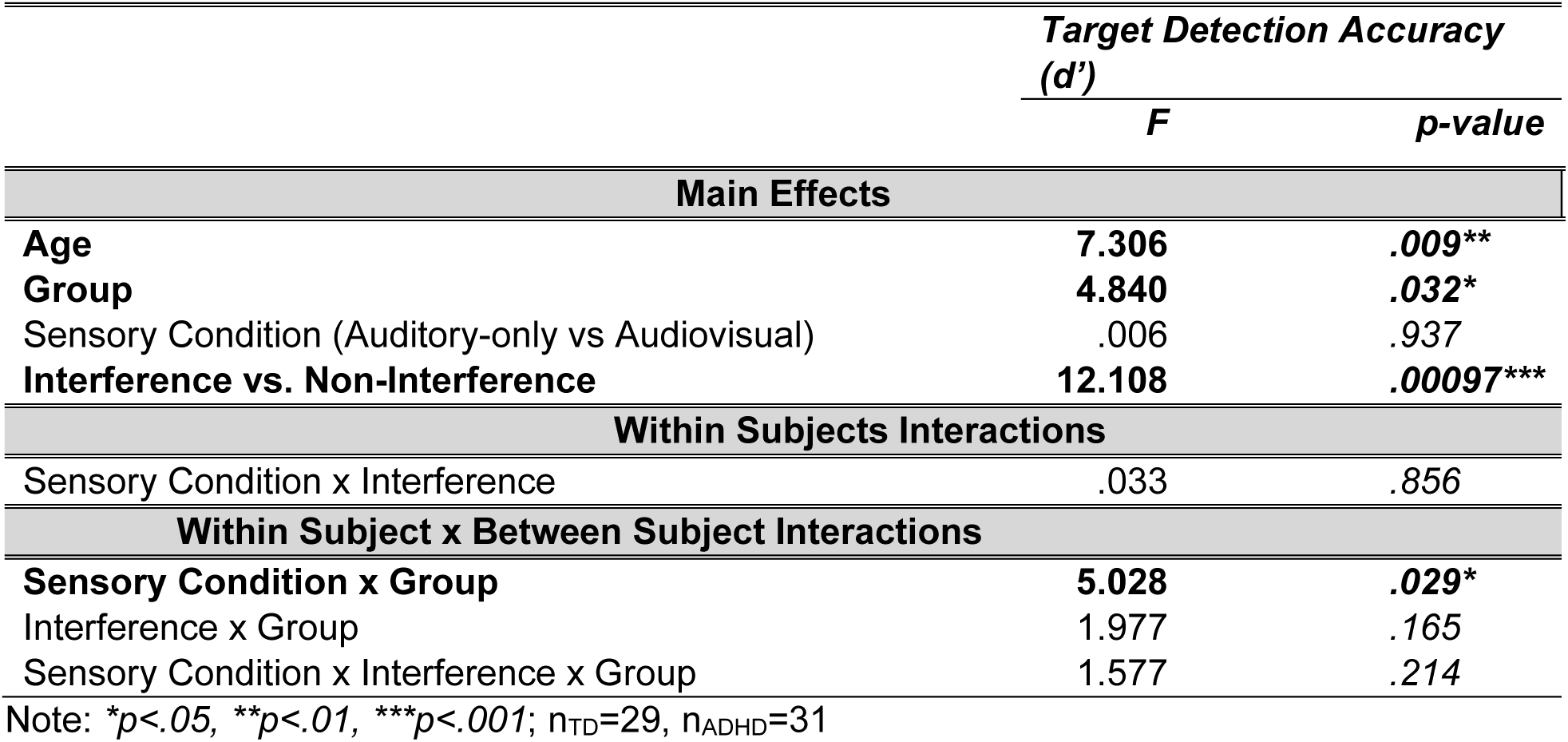
Behavioral Performance.

**Figure 2.**
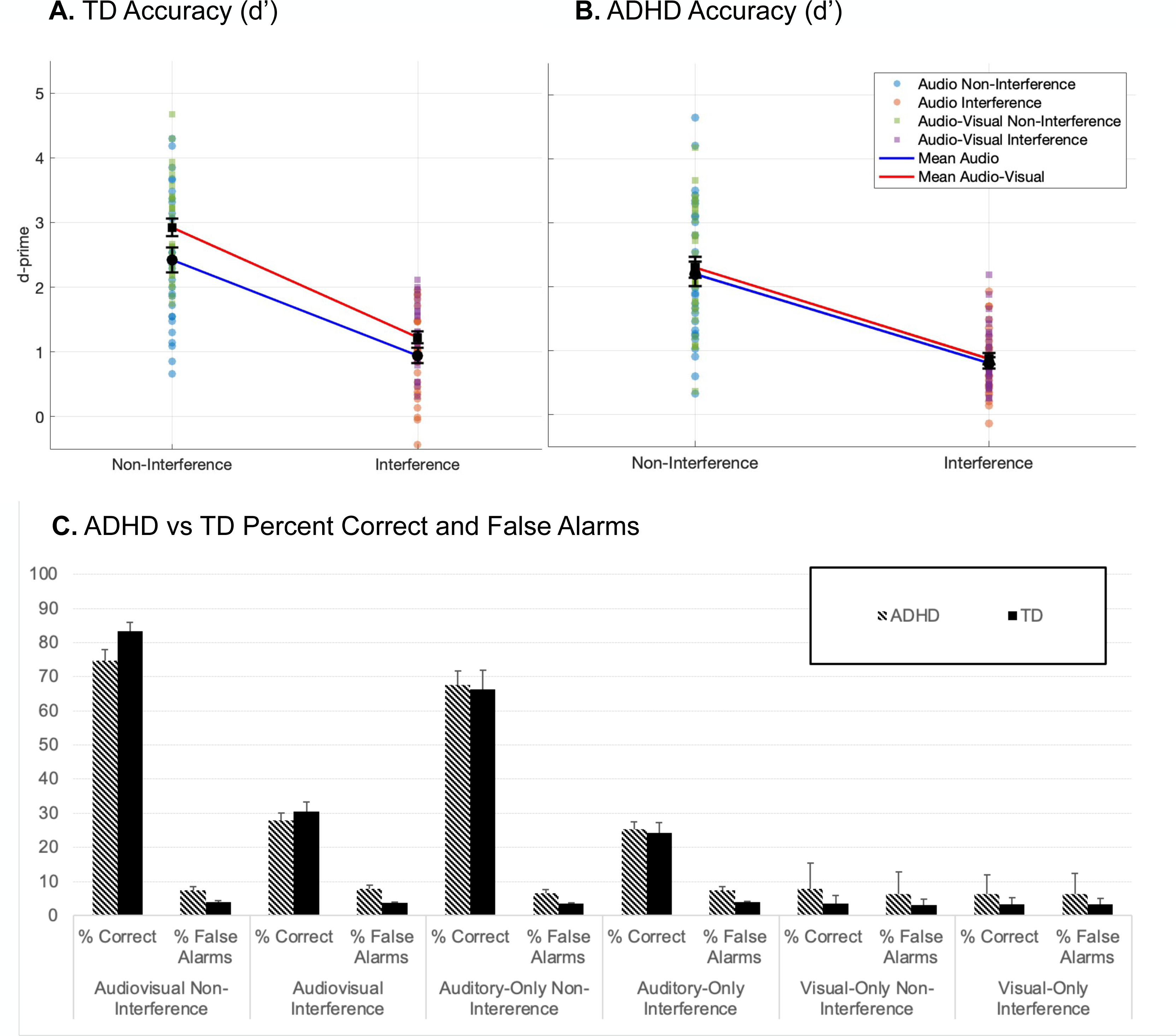
Behavioral accuracy (d-prime) in detecting target speech for children with **(A)** TD and **(B)** ADHD. Group means for the auditory and AV conditions are depicted by the solid blue and red lines, respectively. Individual subject data is represented for the non-interference (auditory-light blue dots; AV-green squares) and interference (auditory-orange dots; AV-purple squares) conditions. **C)** Comparison of behavioral accuracy (percent correct and false alarm responses) between the ADHD (striped) and TD (solid) groups for each of the experimental conditions. Error bars represent the standard error of the mean.

**Figure 3.**
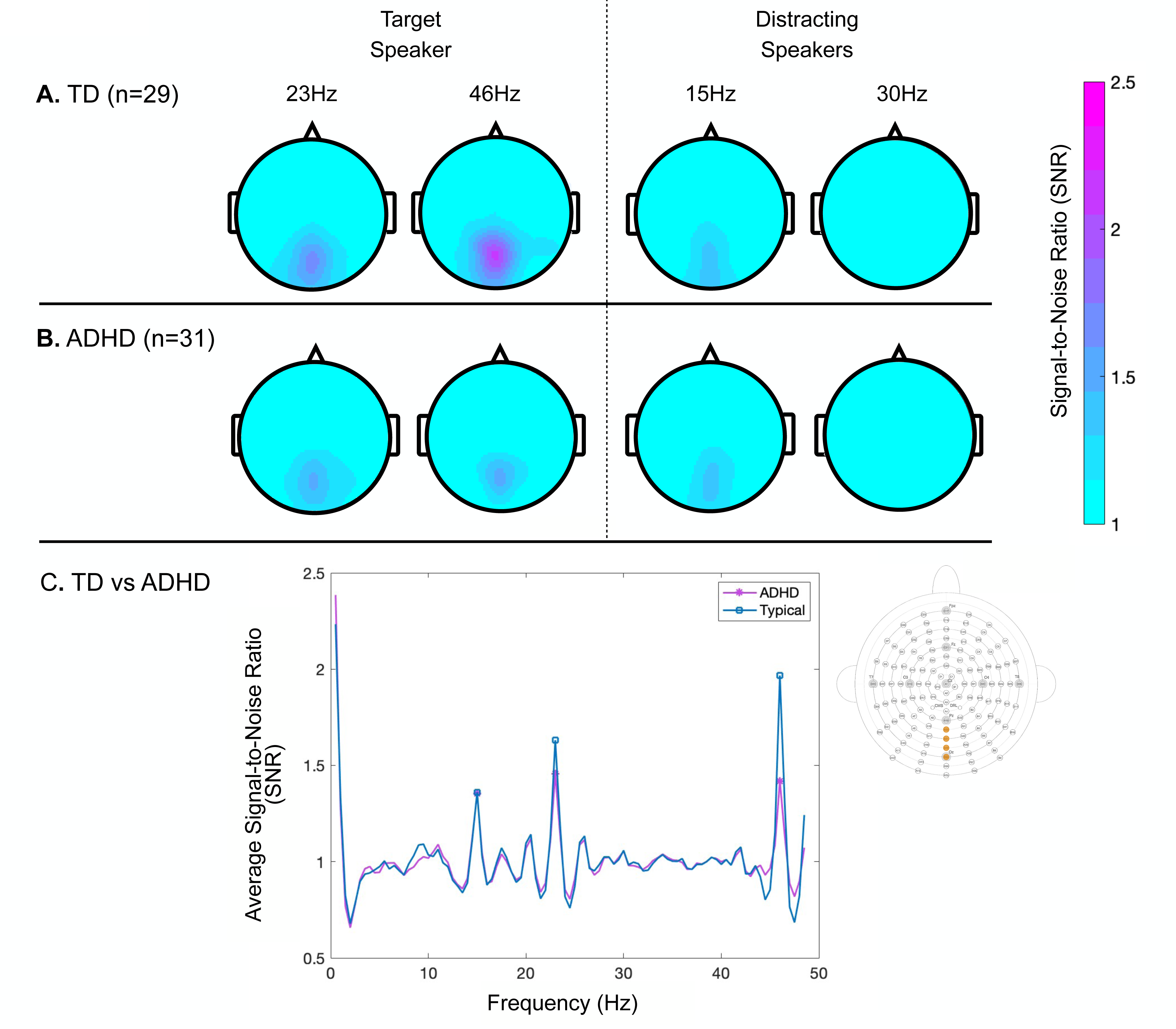
Topographic depiction of steady state visual evoked potentials (SSVEP) in the frequency band of the target speaker (left panel, 23Hz and 46Hz) and distracting speakers (bottom panel, 15Hz and 30Hz for the **(A)** TD and **(B)** ADHD. The color scale depicts the signal-to-noise ratio (SNR) of SSVEP across all 128 scalp electrodes. **C)** Average signal-to-noise SNR amplitude spectra across the four electrodes of interest in the midline occipital to occipital-parietal region for the TD (blue) and ADHD (purple) groups. Asterisks mark the significant spectral peaks at the stimulating frequencies (first harmonics-15 Hz (distractor) and 23 Hz (target); second harmonic-46 Hz (target).

### Steady State Visual Evoked Potentials

Topographic and amplitude spectrum representations of SNR for each of the target and distractor frequencies are depicted by group in Figure 3. There was a main effect of age whereby SSVEP increased with increasing age (F(1,57)=5.103, p=.028, η^2^_p_ =.082), though this maturational pattern did not vary between groups. After covarying for age, there was a main effect of group whereby children with ADHD demonstrated an overall reduction in SSVEP compared to TD children (F(1,57)=7.813, p=.007, η^2^_p_ =.121). There was also a group x frequency interaction (F(1,57)=9.168, p=.004, η^2^_p_ =.139). Children with TD had a significantly larger SSVEP to the target than to the distractors, whereas children with ADHD demonstrated a significant reduction in target SSVEP relative to TD controls. As a result, the ADHD group did not present with a significant difference in SSVEP response between the target and the distractor frequencies.

**Table 3.**
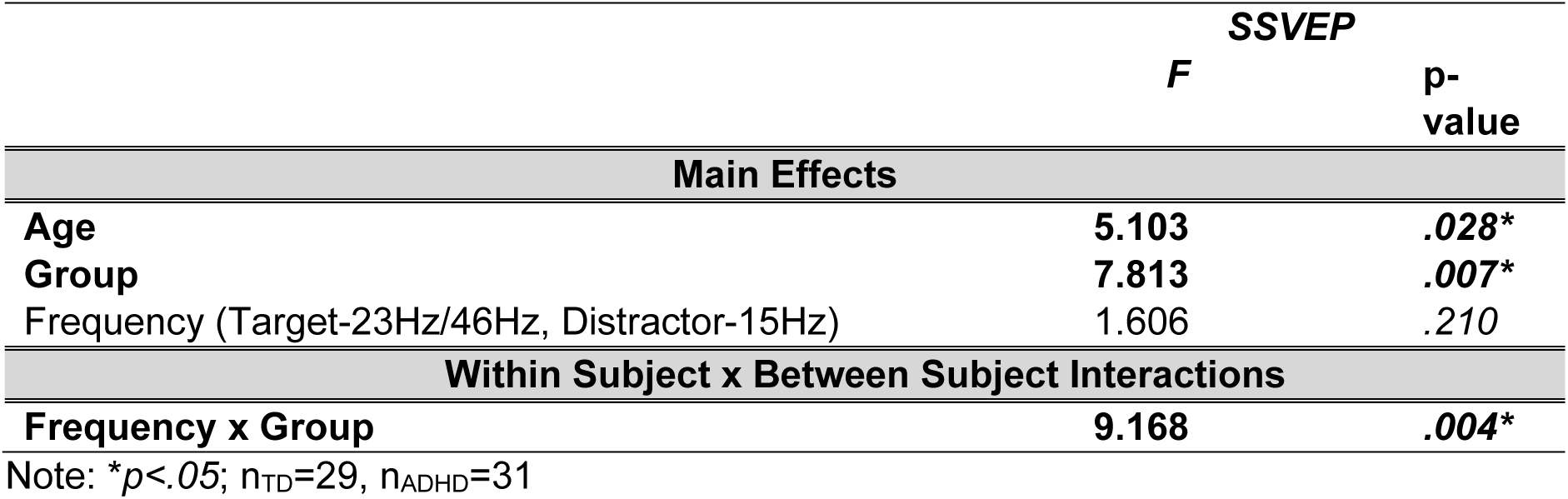
Primary Analysis: SSVEP Signal-to-Noise Ratio (23Hz/46Hz vs 15Hz)

### Neural-Behavioral Correlations

Planned linear regression analyses were conducted separately within each group to explore whether individual variability in target and distractor SSVEP magnitudes predicted target detection accuracy (d’) in each of the AV conditions (interference and non-interference), as visuospatial attention might be expected to influence performance in the AV conditions. Within the TD group, target and distractor SSVEP explained 26.9% of the variance in behavioral performance during the AV interference (F(2,26)=5.844, p=.008, adjusted R^2^=.257) but not AV non-interference (F(2,26)=2.813, p=.078, adjusted R^2^=.115) conditions. This appeared to be primarily driven by a positive relationship between the target SSVEP and accuracy (See Figure 4). SSVEP was not significantly associated with audiovisual behavioral performance in the ADHD group (Interference: F(2,28)=.859, p=.434, adjusted R^2^= -.009; Non-interference: F(2,28)=.006, p=.994, adjusted R^2^= -.071).

**Figure 4.**
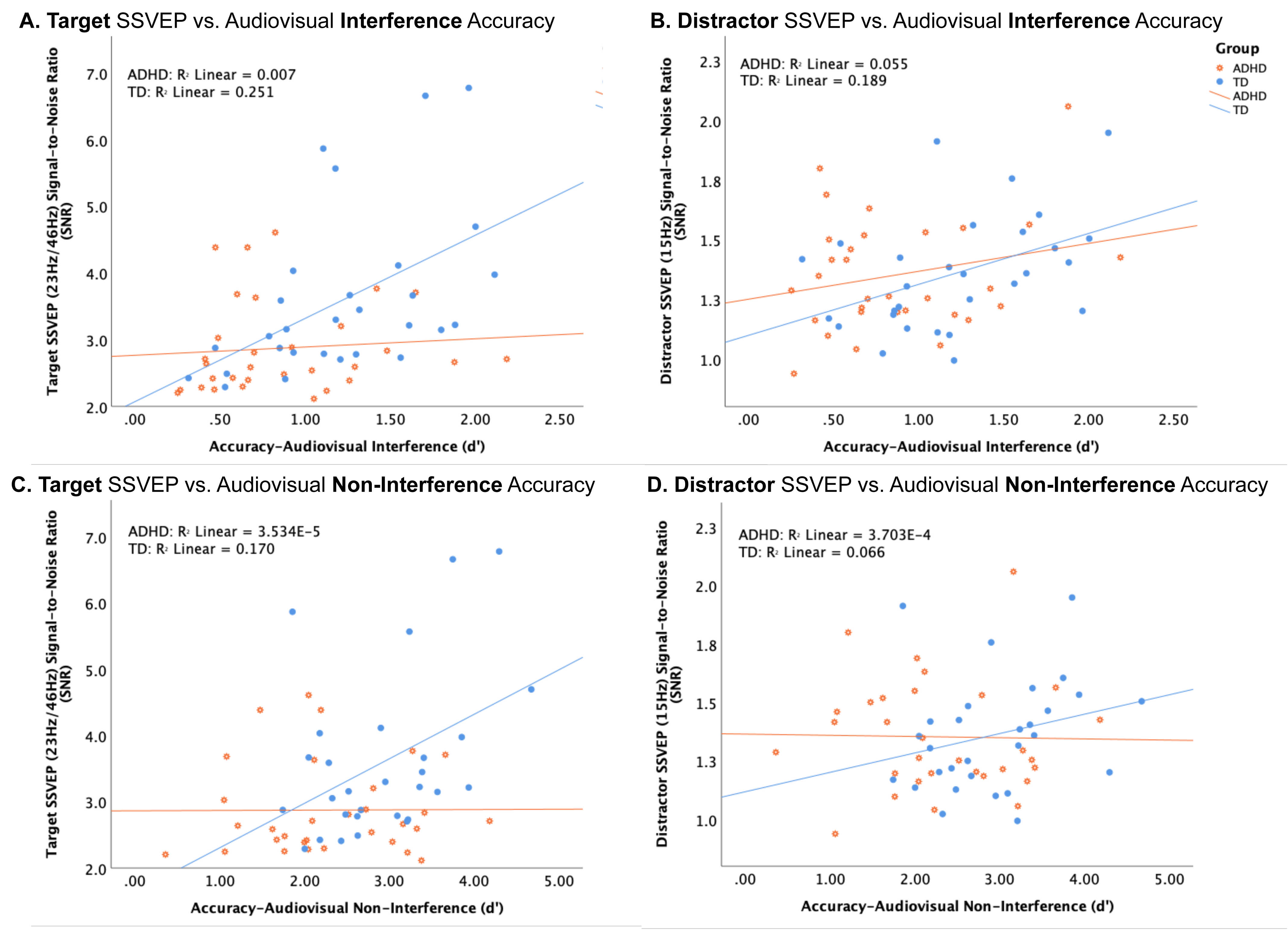
Scatter plots depicting the correlations between audiovisual target detection accuracy and steady state visual evoked potentials (SSVEP) in the TD (blue) and ADHD (orange) groups. **A)** Correlation between target detection accuracy (d’) in the audiovisual interference condition and target (23Hz/46Hz) SSVEP signal-to-noise ratio (SNR). **B)** Correlation between target detection accuracy (d’) in the audiovisual interference condition and distractor (15Hz) SSVEP SNR. **C)** Correlation between target detection accuracy (d’) in the audiovisual non-interference condition and target (23Hz/46Hz) SSVEP SNR. **D)** Correlation between target detection accuracy (d’) in the audiovisual non-interference condition and distractor (15Hz) SSVEP SNR.

Post-hoc exploratory correlational analyses were conducted on the full dataset (inclusive of both diagnostic groups to preserve power and reduce the number of exploratory comparisons) to probe potential relationships between behavioral performance on the VR-EEG speech-in-noise paradigm and performance on the SCAN-3:C Test for Auditory Processing Disorders in Children (See Figure 5). There was no relationship between target detection accuracy on the laboratory paradigm and overall auditory processing composite scores. However, there were positive correlations between target detection accuracy within both the AV and audio-only modalities on the laboratory paradigm and scores on the competing sentences subtest of the SCAN-3:C. These correlations survived correction for multiple comparisons only in the audiovisual interference and non-interference conditions (see Table 4).

**Table 4.** Exploratory Correlational Analyses: Speech-in-noise behavioral performance vs. SCAN-3:C Test for Auditory Processing Disorders in Children.

| Variable |  | Auditory<br>Processing<br>Composite | Auditory<br>Figure<br>Ground<br>(+8)<br>Scaled<br>Score | Filtered<br>Words<br>Scaled<br>Score | Competing<br>Words-Directed<br>Ear Scaled<br>Score | Competing<br>Sentences<br>Scaled<br>Score |
| --- | --- | --- | --- | --- | --- | --- |
| Accuracy Audio<br>Interference (d') | Pearson's r | 0.070 | -0.013 | -0.215 | 0.023 | 0.350 |
|  | *p-value* | 0.631 | 0.928 | 0.134 | 0.873 | 0.013* |
|  | *p-value*<br>*(Bonferroni*<br>*corrected)* | -- | -- | -- | -- | 0.117 |
| Accuracy Audio<br>Non-Interference<br>(d') | Pearson's r | -0.42 | -0.029 | -0.222 | 0.021 | 0.315 |
|  | *p-value* | 0.774 | 0.842 | 0.121 | 0.883 | 0.026* |
|  | *p-value*<br>*(Bonferroni*<br>*corrected)* | -- | -- | -- | -- | 0.234 |
| Accuracy<br>Audiovisual<br>Interference (d') | Pearson's r | -0.234 | 0.144 | -0.211 | 0.146 | 0.440 |
|  | *p-value* | 0.103 | 0.318 | 0.142 | 0.313 | 0.001** |
|  | *p-value*<br>*(Bonferroni*<br>*corrected)* | -- | -- | -- | -- | 0.009* |
| Accuracy<br>Audiovisual Non-Interference (d') | Pearson's r | 0.160 | 0.003 | -0.210 | 0.112 | 0.450 |
|  | *p-value* | 0.267 | 0.984 | 0.143 | 0.438 | 0.001** |
|  | *p-value*<br>*(Bonferroni*<br>*corrected)* | -- | -- | -- | -- | 0.009** |

Note: \* $p < .05$ , \*\* $p < .01$ ; $n_{TD} = 26$ , $n_{ADHD} = 24$

**Figure 5.**
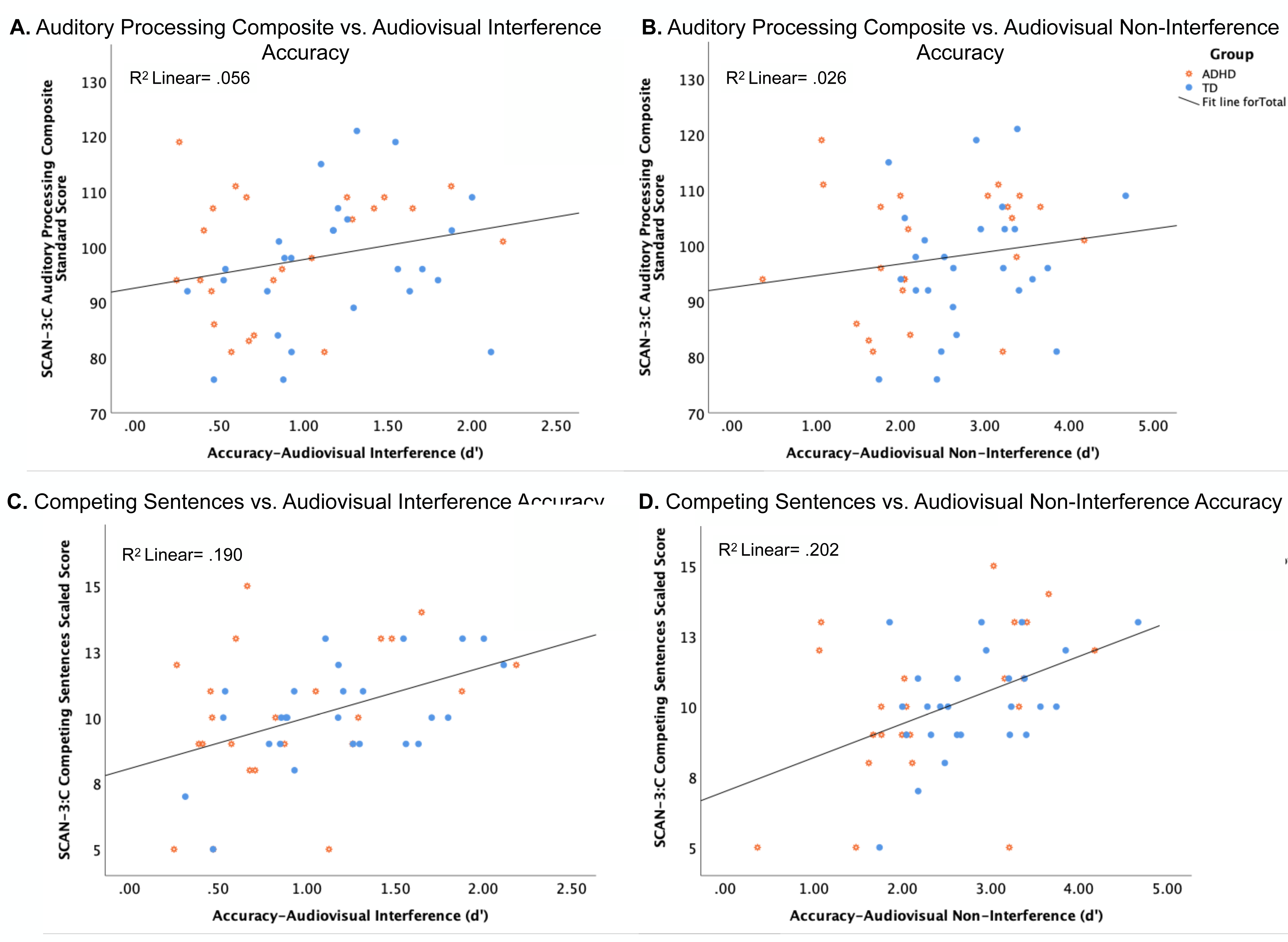
Scatter plots depicting the correlations between SCAN-3:C Tests for Auditory Processing Disorders and audiovisual target detection accuracy in the TD (blue) and ADHD (orange) groups. **A)** Correlations between SCAN-3:C Tests for Auditory Processing Disorders Composite Standard Scores and target detection accuracy (d’) in the audiovisual interference condition. **B)** Correlations between SCAN-3:C Tests for Auditory Processing Disorders Composite Standard Scores and target detection accuracy (d’) in the audiovisual non-interference condition. **C)** Correlations between SCAN-3:C Competing Sentences Scaled Scores and target detection accuracy (d’) in the audiovisual interference condition. **D)** Correlations between SCAN-3:C Competing Sentences Scaled Scores and target detection accuracy (d’) in the audiovisual non-interference condition.

## DISCUSSION

Children with ADHD showed significantly reduced AV gain in both the interference and non-interference conditions. This finding indicates that children with ADHD in the study are not receiving the same benefit of multisensory over unisensory stimuli that TD children exhibit when interpreting speech in the presence and absence of multi-talker noise. This finding is consistent with prior literature showing that adults with ADHD did not improve their understanding of speech in noisy environments when speakers’ faces were visible, unlike the control group, who benefited from the additional visual information (Schulze et al., 2021). Likewise, studies have found that children with ADHD have impaired audiovisual integration, as evidenced by a smaller reduction in inhibition of return for AV targets compared to visual targets in children with ADHD than in TD children (Zhang et al., 2024). Another study investigating early visual and auditory processes in autism spectrum disorder and ADHD found that both groups showed diminished AV interactions compared to TD children (Norcia et al., 2021). However, neither of these prior studies in childhood specifically examined the AV integration of naturalistic lip movements and speech stimuli. Taken together with the existing literature, this study provides compelling evidence for the presence of AV integration deficits sufficient to impact speech perception in ADHD.

To explore whether altered visuospatial attention may contribute to this reduction in AV benefit, we analyzed patterns of visuospatial attention allocation to the three competing speakers as indexed by SSVEP. Children with TD demonstrated robust activation to the target speakers with less activation to the distractor speakers. Children with ADHD showed an overall reduction in the intensity of the SSVEP relative to TD children. However, post hoc tests revealed that this reduction in SSVEP was statistically significant only in the case of the target speaker. As a result, the children with ADHD showed more equivalent activation to the two competing speakers in contrast to the selective enhancement of SSVEP to the target speaker exhibited by the TD children. This finding aligns well with prior literature showing that the subset of TD adults with the highest behavioral distraction levels (as evidenced by false alarm rates) during an AV speech-in-noise task exhibited more equivalent SSVEP between the target and distractor frequency bands (Senkowski et al., 2008). However, unique to the findings in the present investigation, the difference between groups was most pronounced in the intensity of response to the target. In contrast, in the study of TD adults by Senkowski and colleagues, distracted subjects showed augmentation of response to the distractor stimuli (Senkowski et al., 2008). Distinct SSVEP patterns have also been observed in adolescents with ADHD, accompanied by significantly poorer performance in stop-signal tasks involving both visual and auditory stimuli compared to TD controls (Khaleghi et al., 2019). Adolescent males with ADHD also showed reduced SSVEP modulation during the continuous performance task, a widely used neuropsychological measure of sustained attention (Silberstein et al., 1998). Neither of these prior studies employing SSVEP in ADHD explored audiovisual integration; to our knowledge, this represents the first investigation to do so. Nevertheless, there is agreement across studies in the finding of disrupted SSVEP correlating with performance on cognitive tasks across a variety of stimuli in both children and adolescents with ADHD.

As outlined in the introduction, there is a compelling body of work supporting the sensitivity of SSVEP to visuospatial attentional allocation. Thus, the reduction in SSVEP we observed in children with ADHD implies disrupted attentional modulation to visual information during audiovisual speech processing. This notion is in agreement with many prior studies. Children with ADHD tend to have poorer accuracy in visual selective attention and visual search performance tasks (Hokken et al., 2023). They also spend less time viewing relevant areas of images and take longer to respond when detecting emotions compared to children without ADHD (Serrano et al., 2018). Though that study did not focus on lip movements, it suggests a broader pattern of altered visual attention in ADHD that could potentially affect the processing of visual speech cues, consistent with the observations in the present study.

Within the TD group in our sample, there was a positive relationship between the strength of the SSVEP to the target speaker and the amount of behavioral AV gain during speech-in-noise perception, implying that a reduction in visuospatial attention directed to the target speaker may underlie lower accuracy in speech detection in multi-talker scenarios. However, it is important to note that a causal relationship can be inferred based on the study design. Interestingly, this neural-behavioral relationship was not observed within the ADHD group. A weaker relationship between SSVEP and AV gain in children with ADHD may imply an additional influence of other unmeasured factors beyond visual attention that contribute to the marked attenuation of AV gain in these children. This study did not measure participants’ ability to sustain *auditory* (only visual) attention during the VR-EEG speech-in-noise task. However, in exploratory analyses, behavioral performance on this task did appear to correlate with dichotic listening to competing sentences presented in an auditory-only format. Impaired performance on dichotic listening and sound localization (Fu et al., 2022) has been described in ADHD previously (Manassis et al., 2000) and may imply a role for selective auditory attention in addition to visual attention contributing to AV speech perception in noisy settings. This is perhaps unsurprising given the well described cross-modal influence of many attentional inputs (Teder-Salejarvi et al., 1999; Zhao et al., 2024).

Additionally, though outside the scope of this investigation, these findings raise an interesting question of whether the differences in visual attention among children with ADHD are specific to the speech domain as tested here or would also extend to non-speech AV or even unisensory visual-only stimuli. Given that the SSVEP in children with ADHD was reduced overall (to both the target and the distractor) relative to controls, the disruption does not appear to be solely in the ability to selectively enhance attention to relevant speakers and suppress attention to irrelevant speakers. Instead, this pattern suggests an overall reduction in the maintenance of visual attention to both types of stimuli and/or a deficit in visual entrainment to oscillatory stimuli. Prior research has found that many children with ADHD have higher SSVEP amplitudes, suggesting that, at a minimum, there is no specific deficit in the ability to generate an SSVEP in ADHD (Khaleghi et al., 2019). One possible explanation of our findings then is that with the simultaneous presentation of visual and auditory stimuli, children with developmental disabilities may shift their attention away from the visual modality to process the auditory stimuli. Consistent with this possibility, prior studies have shown a decrease in visual activity when attention is shifted from vision to audition (Foxe et al., 1998; Foxe et al., 2005; Shomstein and Yantis, 2004).

## LIMITATIONS

This study has clear implications for central auditory processing disorder, a condition characterized by difficulty interpreting speech, especially in noisy settings, despite normal hearing thresholds. However, we did not require that children with ADHD be additionally diagnosed with auditory processing disorder as part of the inclusion criteria, seeking instead to evaluate mechanisms of speech-in-noise processing across ADHD in general. Given the high degree of overlap in symptomatology, there is marked difficulty in accurately diagnosing central auditory processing disorder in children with inattention (Cacace and Enayati, 2022; Moore, 2015). Additionally, many children with ADHD have difficulty in auditory processing, even when their presentation is ultimately deemed subthreshold for confirmation of a central auditory processing disorder. Indeed, children with ADHD in our study demonstrated pronounced deficits in their ability to process speech despite lacking central auditory processing diagnoses and being matched with the TD population on verbal comprehension indices. This elevates the value of examining these symptoms along a continuum.

Lastly, eye-tracking capability was not available at the time of the initial experimental design because the stimuli were presented on the Meta Quest 2. As a result, we are unable to distinguish between disruptions to overt orienting of gaze to relevant stimuli and covert shifting of mental attention to the relevant stimuli without concomitant shift in gaze. Future studies can further delineate the specific types of attentional orienting as newer VR headsets offer in-built eye-tracking capability.

## SUMMARY

Children with ADHD demonstrate a reduction in the typical perceptual benefit afforded by exposure to multisensory (AV) vs. unisensory speech stimuli, accompanied by a substantial difference in the allocation of visuospatial attention to the relevant speaker. Thus, dysregulation of selective visuospatial attention may impact their ability to perceive language in noisy settings effectively.

---

**Figure S1.**
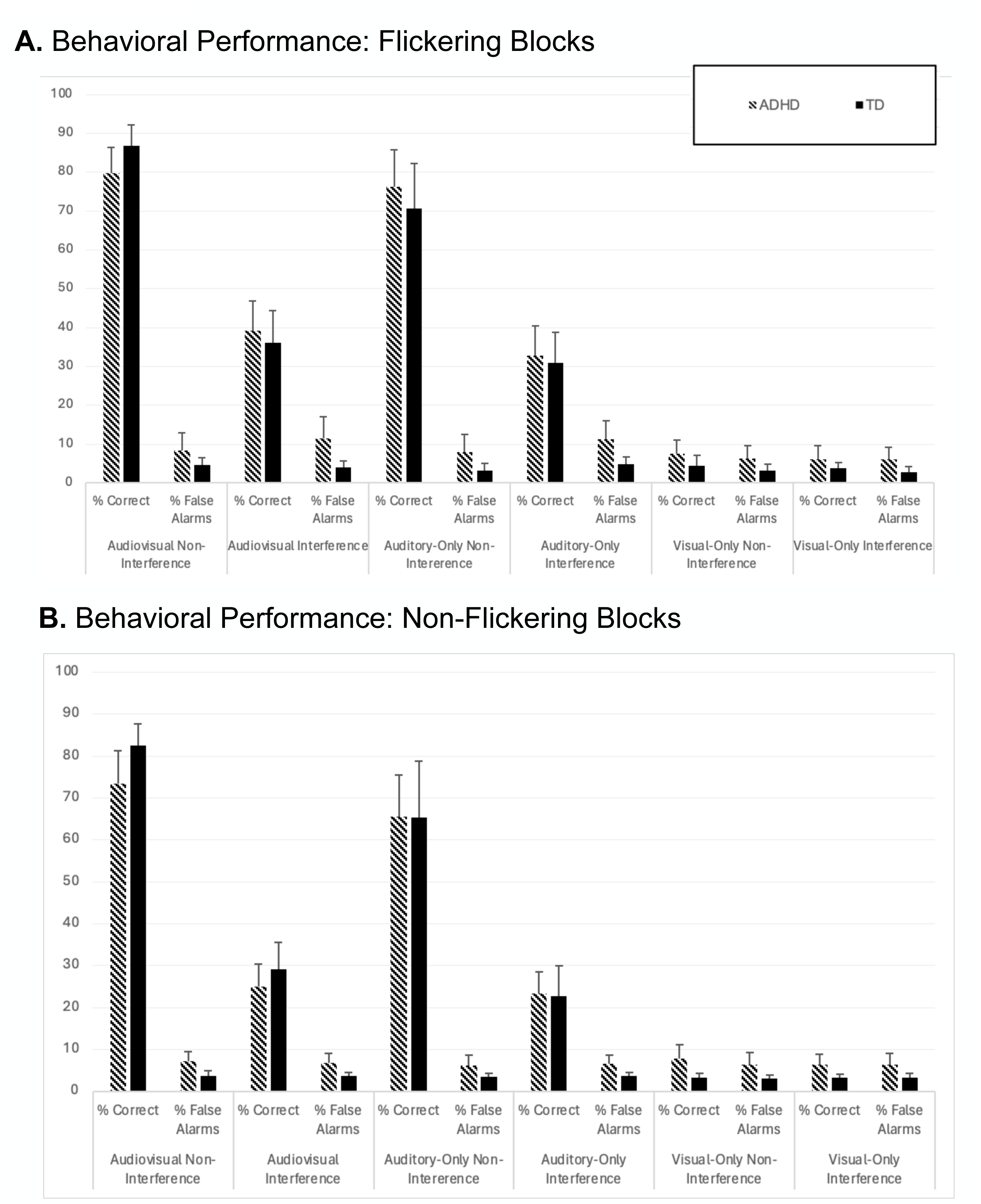
Comparison of behavioral accuracy (percent correct and false alarm responses) between the ADHD (striped) and TD (solid) groups for each of the experimental conditions across the **(A)** flickering and **(B)** non-Flickering experimental blocks. Error bars represent the standard error of the mean.

**Table S1.** SSVEP Signal-to-Noise Ratio (23Hz vs 46Hz vs 15Hz vs 30Hz)

|  | **SSVEP** |  |
| --- | --- | --- |
|  | *F* | *p-value* |
| **Main Effects** |  |  |
| **Age** | **4.870** | **.031\*** |
| **Group** | **7.659** | **.008\*** |
| Frequency (Target 1<sup>st</sup> Harmonic/23Hz, Target 2<sup>nd</sup> Harmonic/46Hz, Distractor 1<sup>st</sup> Harmonic/15Hz) | 0.116 | .842<sup>a</sup> |
| **Within Subject x Between Subject Interactions** |  |  |
| **Frequency x Group** | **8.714** | **.0009<sup>a</sup>** |

Note: \* $p < .05$ ; $n_{TD}=29$ , $n_{ADHD}=31$ ; <sup>a</sup>Greenhouse-Geisser corrected for violation of sphericity

## ACKNOWLEDGMENTS

The authors acknowledge the important contributions of Kimothy Gingrich, a psychometrist, who assisted with a subset of the phenotyping data collection and Dasle Han, a University of Rochester undergraduate research assistant, who assisted with final phenotyping data curation.

## AUTHOR CONTRIBUTIONS

E.J.K and J.J.F conceived and designed the research; N.J.D, S.M, and Z.B performed the electrophysiologic data collection, A.R., R.R., C.L., and E.J.K. conducted and interpreted the neuropsychological testing data. N.J.D. and E.J.K analyzed data; E.J.K, N.D and J.J.F. interpreted results of experiments; N.J.D and E.J.K prepared figures and drafted the manuscript. All authors revised the manuscript and approved the final version of manuscript.

